# Modeling Type-I Fatty Acid Synthase with Acyl Carrier Protein at Ketoacyl Synthase

**DOI:** 10.64898/2026.05.15.725316

**Authors:** Manisha Sharma, Harshwardhan H. Katkar

**Affiliations:** Department of Chemical Engineering, Indian Institute of Technology Kanpur, Kanpur 208016, Uttar Pradesh, India

## Abstract

Mycobacterium tuberculosis fatty acid synthase I (Mtb FAS-I) is a multifunctional hexameric complex essential for fatty acid (FA) synthesis. The need of a hexameric structure for activity of the complex in Mtb remains elusive. Here, we model a conformation of the functionally active complex with acyl carrier protein (ACP) at ketoacyl synthase (KS). Our model reveals a crucial cross-dome dependence in the mechanism of FA synthesis at the condensation step. Using molecular dynamics simulation, we identify key ACP and KS residues that mantain persistent interactions. ACP’s phosphopantetheine (PPT) arm adopts several conformations while accessing KS’s catalytic pocket, including two distinct conformations that correlate with volumes of ACP and KS pockets. A PHE residue, reported as a gatekeeper of the KS pocket in other species, also shows open and closed orientations in our simulation. Our results provide crucial insights that are essential for a mechanistic undersanding of the Mtb FAS-I complex.

## 1 INTRODUCTION

The reaction mechanism of fatty acid synthesis has long been established and is conserved across several species, including yeast, bacteria, and mammals. Fatty acid synthase (FAS) type I and type II are a class of proteins involved in fatty acid production. FAS-I complexes are structured proteins that form either a hexamer in bacteria or a homododecamer in fungi with all catalytic enzymes embedded in a single or two subunits, respectively. In contrast, these same components exist as independent enzymes in the FAS-II system. Previous studies on yeast and bacterial fatty acid synthase type I (FAS-I) have shown that hexameric assembly is necessary for an active FAS-I complex [32, 22, 10]. A recent study on mycobacterial species, *Mycobacterium tuberculosis* (Mtb), has also shown that the FAS-I complex is active in hexameric form, only after a post-translational modification of Mtb acyl carrier protein (ACP) with a prosthetic arm, *viz*, phoshopanthethein (PPT) arm [3]. It is quite straightforward why modified ACP is required for fatty acid synthesis, as the PPT arm carries the substrate along with it during the ACP shuttling mechanism. It remains unclear whether the post-translational modification precedes or follows assembly formation, and the mechanism by which ACP becomes encapsulated within the reaction chambers is still unresolved. Notably, successful post-translational modification of ACP has been reported to occur independently of assembly [10]. Furthermore, even when the condensation enzyme in yeast FAS-I is inhibited, yeast ACP has been reported to exhibit partial occupancy across all catalytic domains [12]. Literature suggests that both of these prerequisites must be satisfied for efficient fatty acid synthesis. Together, these observations raise a fundamental question: what governs the successful progression of fatty acid synthesis within this system? If both conditions are essential, then what evolutionary advantage might underlie the hexameric architecture of the Mtb FAS-I complex? All the catalytic domains and the carrier protein, ACP, are arranged in a single monomer, and even the independent catalytic domains (FAS-II enzymes) remain active and carry out catalysis via diffusion in the cell relying on abundance of the ACP molecules [2, 6, 44]. This structural information supporting the need for hexameric assembly in FAS-I has not been reported in the previous literature.

Many structural details are available for the fungal FAS-I complex, whereas only limited information is available for the mycobacterial FAS-I. The lack of information also limits our understanding of the ACP shuttling mechanism. The existing structural details provide valuable insights; however, they do not specifically address how structural bias drives assembly, how the two domes function, or how ACPs communicate within each dome. For nearly three decades, there had been a long-standing debate regarding the fungal FAS-I complex, specifically whether it exhibits half-site or full-site reactivity, cooperativity or lacks cooperativity, and whether the active centres function independently or in a coordinated, interdependent manner [41, 33]. The debate was settled with full site reactivity. The two chambers of fungal FAS-I have been interpreted as two independent domes [2]. What evidence supports the independence of the domes? To date, there have been limited efforts to explore how such questions could be addressed using the structures of FAS complexes. A more comprehensive understanding of the relationship between FAS-I structures and the ACP shuttling mechanism still needs to be established.

In this manuscript, our focus is on the condensation step of the ACP shuttling mechanism. We will discuss how this step is crucial at various depths of understanding FAS-I structure and the fatty acid synthesis. Condensation occurs at the ketoacyl synthase (KS) enzyme, and it plays an important role in controlling the length of resultant fatty acids [14, 38, 39]. The literature is rich with fungal FAS-I structures in which ACP is stalled at the KS domain. Also, in Mtb FAS-I, an electron density of ACP has been reported near the KS domain [9]. KS is reported to have the highest ACP occupancy (≈ 30%), along with acetyl transferase (AT) and enoyl reductace (ER), as ACP visits KS twice per reaction cycle [12]. Why is this interaction of ACP at KS intrinsically favoured? A reason cited is that the equatorial KS domain is least affected by changes in the complex’s shape due to large domain movements of the barrel wall [12]. The KS domain assembles into three dimers within the wheel region, each formed by pairing a KS domain from the upper and a lower dome. This represents a key structural feature of the Mtb FAS-I complex and is conserved across species. In fungal FAS-I, the corresponding dimer is formed between two distinct *α*-subunits containing the KS, ketoacyl reductase (KR), PPT, and ACP domains, whereas in Mtb FAS-I, the same dimeric arrangement arises from two polypeptides and each harbours all catalytic domains. The steps involved in the assembly formation of the *α*_6_ *β*_6_ complex in fungal FAS-I have recently been investigated [10]. However, Mtb FAS-I assembly formation of six identical polypeptides has not been reported yet. It has been reported that KS dimerisation is crucial for fungal FAS-I assembly formation and the post-translational modification of ACP at the PPT domain. On the other hand, the PPT domain is absent in Mtb FAS-I. Thus, assuming the same for Mtb FAS-I would be inaccurate. What is the advantage of the KS dimer formation in Mtb FAS-I? Notebly, the KS domains of each dome face in different reaction chambers [26, 16]. Then the question rises, how does the ACP carry out the condensation reaction if its own KS catalytic site faces a different dome?

Recent reported structures of fungal FAS-I resolved ACP at different domains, namely AT, KR, ER, and malonyl/palmitoyl transferase (MPT), with SER180 residue of ACP recognition helix (*α*-2) pointing towards their respective catalytic clefts with the PPT arm within catalytic range of 18 Å (approximate length of PPT arm) from the catalytic residues [18, 40, 29, 37]. In contrast, the PPT arm has not been resolved within the KS pocket. The PPT arm is involved in the growing fatty acid chains by accessing the catalytic residues. It has been proposed that the condensation of acyl derivatives is in general limited by how the substrate reaches the vicinity of catalytic residues [13]. The PPT arm has been hypothesised to have a switch-blade mechanism and to be dynamic in nature [26]. It adopts a different conformation to reach the catalytic site and carry out catalysis [16].The electron density of the PPT arm inside the KS pocket in yeast FAS-I has been observed, and a model was proposed with the PPT arm inside the pocket [26]. No further information is available regarding the conformation of PPT arm in the KS domain. Also, the condensation reaction mechanism suggests that the substrate acetate or malonyl must be in close contact for bond formation, and the PPT arm facilitates that, which is approximately 18 Å-21 Å in length [37]. Then how does the PPT arm come into the range of catalysis? As ACP and PPT arm have been highly conserved across different fatty acid machineries such as FAS-I, FAS-II and polyketide synthase (PKS) systems and across different species. The PPT arm dynamics can serve as a probe for tracking fatty acid synthesis [19]. The flexible nature of the PPT arm and the transient protein-protein surface interactions between ACP and its catalytic domains pose a major challenge for understanding whether ACP carries out fatty acid synthesis by random diffusion or along a prescribed pathway. Localizing and characterizing the PPT arm in experiments is highly challenging due to its fast dynamics [42].

In this work, we provide structural and mechanistic insights into the need of a hexameric structure of the Mtb FAS-I complex and its implications on ACP’s shuttling mechanism. In this context, we identify a unique cross-dome binding mode of ACP that arises from intrinsic structural constraints within the assembly. To achieve a functionally coherent framework, we model the structure of the previously unresolved ACP, and incorporated all relevant cofactors and tethering linkers, thereby constructing the first complete structural model of a catalytically competent FAS-I complex in a conformation with ACP at KS. The model provides direct structural evidence for the need of a hexameric structure for an active complex. Molecular dynamics simulation of our model helps us characterize persistent residue-level contacts between ACP and KS. We also characterize conformational dynamics of the PPT arm and investigate two key sets of conformations: one with the PPT arm inside the KS pocket and another with it sequestered in ACP. We report a correlation between PPT arm conformations and sizes of these pockets. We also report the dynamics of a PHE residue in the KS pocket, which has been previously identified as a gating residue in the KS pocket. Lastly, we present structural insights into ACP’s accessibility to the three KS domains within the same dome.

## 2. Methods

### 2.1 Model Development

The cryogenic electron microscopy (cryoEM) structure and FASTA sequence of Mtb FAS-I (PDB-ID 6GJC) homo-hexamer, with missing regions corresponding to ACP and its tethering linkers, were retrieved from the Protein Data Bank. [9] The cryoEM structure is shown in Figure 1 (a). The missing region corresponding to ACP in a single chain (residues 1756-1925) was modelled using the *segment modeling* feature of MODELLER 10.4, with homologous ACP structures of *S.cerevisiae* (SC) (PDB ID 8PS1) and *Candida albicans* (CA) (PDB ID 6U5V) as inputs. [40, 28] Model with the lowest dope score was selected as our ACP model. The structure in PDB ID 8PS1 was moved in order to superimpose its KS region with the corresponding region of chain A of PDB ID 6GJC using Pymol, followed by superimposing our ACP model on its ACP region, to obtain a conformation with ACP at KS.

**Figure 1.**
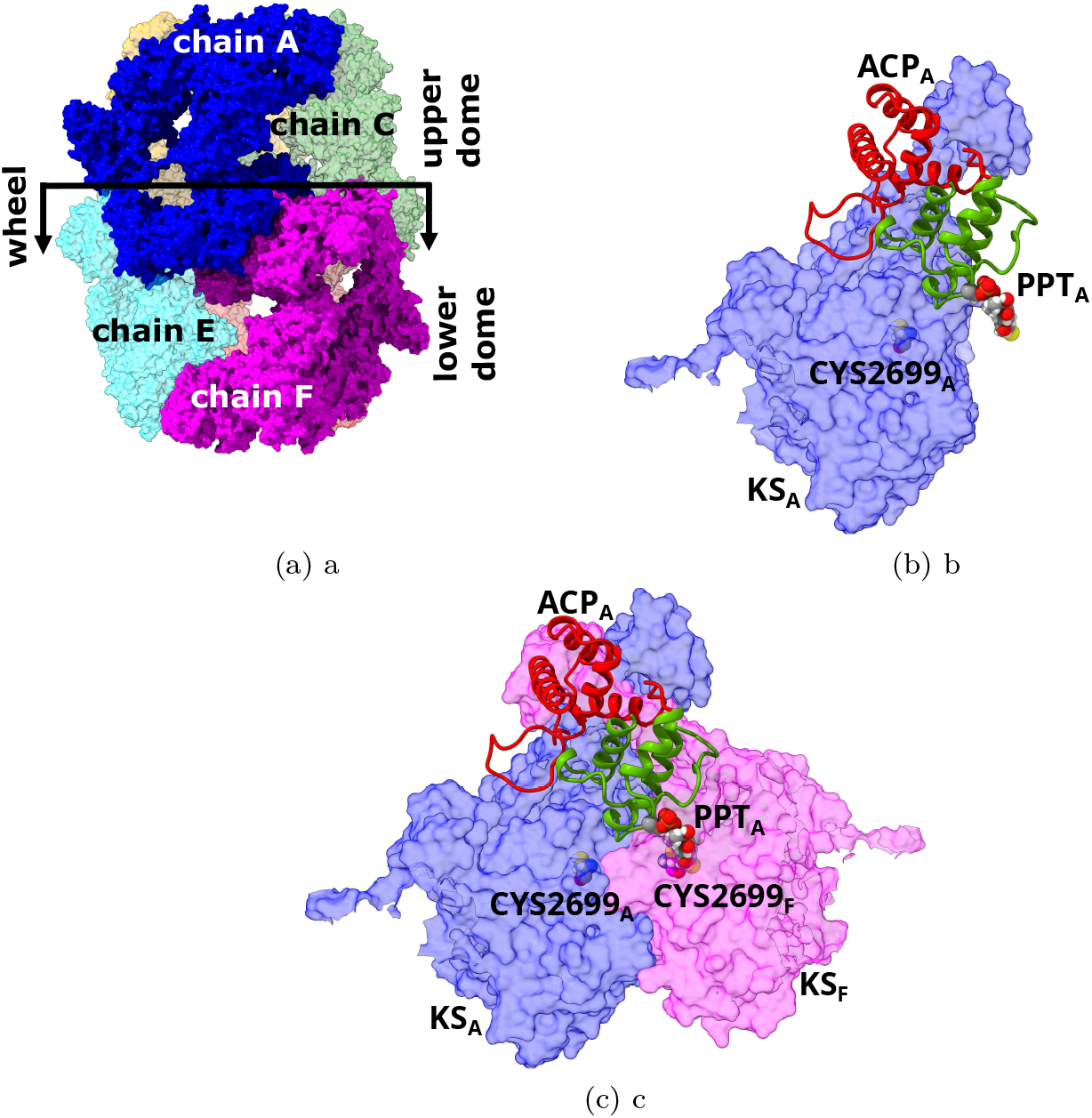
Mtb FAS-I structure. (a) The hexameric complex (PDB ID 6GJC [9]). Different colors represent different chains. (b) Modeled conformation of ACP_A_ (ribbon representation) at KS_A_ (surface repesentation). The PPT arm and CYS2699 atoms are shown as balls. (c) The same conformation as (b) with additional lower dome KS_F_ shown.

The resulting ACP model showed good agreement with recently resolved Mtb FAS-I ACP structure (PDB ID 9PQY), models predicted by TASSER web server and AlphaFold web server, and our earlier model. [39, 11, 46, 24]. The root mean square deviation (RMSD) based on C_*α*_ of all ACP residues (1755 to 1925) of these structures relative to our ACP model was 4.688 Å, 3.344 Å, 4.538 Å and 8.446 Å, respectively. The mismatch was primarily due to the loops present in ACP. The RMSD based on C_*α*_ of helix-forming residues (residues 1755 to 1765, 1778 to 1784, 1786-1802, 1813-1825, 1832 to 1848, 1853 to 1863, 1868 to 1883 and 1904 to 1925) reduced to 2.943 Å, 2.867 Å, 2.849 Å and 7.603 Å, respectively (see Figure SI1). Our earlier model included only a part of the helix *α* − 7, which is refined in our current model. [24] The rest of the helices are nearly identical between our earlier and current model.

Missing regions, *viz*. the tethering linkers PL (residues 1710-1751) and CL (residues 1926-1946) along with other unstructured residues (2254-2261, 2302-2307, and 3015) were modelled using the *loop modelling* feature of Modeller 10.4. This resulted into a complete model of a single Mtb FAS-I chain with ACP at KS, albeit with missing cofactors. The cofactors FMN and NADP in ER of this chain were modeled by superimposing ER of PDB ID 8PS2 with the corresponding region of chain A of PDB ID 6GJC, while the cofactor NADP in KR of the chain was modelled by superimposing KR of PDB ID 8PRV with the corresponding region of chain A of PDB ID 6GJC.

Next, the residue SER1787 was modified to include the post-translationally tethered PPT arm. The structure of recently resolved human FAS-I with PPT arm-attached ACP at KS (PDB ID 9MJ9) was used as a template. [7] KS from PDB ID 9MJ9 was aligned to the corresponding region of chain A of PDB ID 6GJC to obtain an initial conformation of the PPT arm. The Pymol *bond* command was used to create a bond between the atoms OG of SER1787 and P of the PPT arm. The conformation of PPT arm was slightly refined by ~ 1 Å using the Pymol command *translate [0,1,0]*.

The resulting complete single chain of Mtb FAS-I was inclusive of all cofactors and the post-translational PPT arm modification. Post-minimization (details below), the chain was super-imposed with each chain of PDB ID 6GJC to obtain a model of homohexameric Mtb FAS-I complex with ACP of each chain stalled at its KS. We label this specific conformation as KS’.

A similar process was followed to generate two additional ACP conformations, KS” and KS”‘. In these conformations, ACP of each chain is stalled at a KS at one of its two neighboring chains in the same dome. Labeling the six chains as A,B, C (upper dome in Figure 1 (a)), D,E and F (lower dome), and using subscripts to denote the chain of a given domain, the three conformations are characterized as follows. KS’ denotes a conformation where ACP_*A*_ stalled at KS_*A*_, KS” denotes ACP_*A*_ stalled at *KS*_*B*_, and KS”‘ denotes ACP_*A*_ stalled at *KS*_*C*_. All remaining chains are in an identical conformation, with their ACP stalled at KS of respective chains.

### 2.2 Simulation Details

The CHARMM-GUI Input Generator Solution Builder tool was used to prepare all simulation systems. [17] CHARMM36 force-field was used for all standard residues, ions and TIP3 water. [21, 5, 15, 25] The topology and parameters for cofactors FMN, NADP and the PPT arm were generated using CHARMM General Force Field (CGenFF). [43, 45] GROMACS 2024.4 was used for performing simulation. [1]

A single chain of Mtb FAS-I with cofactors and PPT arm was neutralized with NaCl and solvated using the TIP3 water model. The system was energy minimized, followed by 3 ns simulation with restraints on all protein atoms minus PL, using NVT ensemble at 310 K implemented using V-rescale thermostat to generate an equilibrated conformation of the single chain without overlaps. This conformation was used to generate the hexameric complex.

The complex was neutralized and solvated, resulting into a simulation box with ~3.2 million atoms. Energy minimization was performed with restraints of 500 kJ mol^−1^ nm^−2^ on backbone atoms and 100 kJ mol^−1^ nm^−2^ on side chain atoms of the protein using the steepest descent algorithm. The system was gradually heated from 0 K to 310 K in 5 ns and then maintained at 310 K for additional 5 ns with restrains of 1000 kJ mol^−1^ nm^−2^ on all protein, co-factor and PPT atoms using NVT ensemble with V-rescale thermostat. Restraints were relaxed stepwise over the next 106 ns using the NPT ensemble at 1 bar pressure and 310 K temperature, implemented using C-rescale barostat and V-rescale thermostat. A restraint of 1000 kJ mol^−1^ nm^−2^ was applied on all protein, cofactor and PPT atoms for the first 10 ns, followed by successively relaxing the restraints on (a) all hydrogen atoms for 20 ns, (b) all side chains for 20 ns. The restraint on the protein backbone minus ACP, PL and CL was reduced stepwise from 1000 kJ mol^−1^ nm^−2^ to 1 kJ mol^−1^ nm^−2^ over 56 ns. Unconstrained simulation of the complex was carried out for additional 300 ns to generate a simulation trajectory. Five independent 300 ns-long unconstrained trajectories were generated and used in subsequent analysis.

### 2.3 Pocket Volume Calculations

Pocket Volume Measurer (POVME) 2.0 tool was used to calculate pocket volumes of both, the ACP catalytic core and the KS catalytic pocket. [8] Seven inclusion spheres were identified by trial-and-error to describe the ACP catalytic core pocket. Similarly, two inclusion spheres were used to describe the KS catalytic pocket. Pocket volume for each simulation frame was calculated using a grid spacing of 1.0 Å, and a cutoff distance of 1.09 Å.

## 3 Results and Discussion

### 3.1 Hexameric structure of the active complex: Direct cross-dome interaction between ACP and KS

In the proposed reaction mechanism of FAS-mediated fatty acid synthesis, KS is involved in transacylation and condensation reactions. The former involves transfer of either acetyl moiety or the growing fatty acid moiety that is connected to the S atom of the PPT arm of ACP onto S atom of CYS residue of KS. The latter involves condensation of malonyl moiety onto the transacylated substrate. [30]

Despite the inclusion of all required enzymatic domains in a single chain, Mtb FAS-I requires to assemble into its hexameric complex in order to actively synthesize fatty acids. [32, 9] Structural analysis of our model of the complex with ACP at KS domain provides a crucial insight into the need of this hexameric structure for activity of the complex. In the Mtb FAS-I complex, all KS domains along with all KR domains from the six chains form the wheel region, which structurally partitions the complex into an upper dome and a lower dome (Figure 1a). The three ACP domains of one dome are sterically restricted from crossing the wheel to enter the opposite dome. This compartmentalization of ACP has been interpreted to suggest that the fatty acid synthesis follows a mechanism in which the two domes work independently. [2] However, interpreting the domes as functionally independent can be misleading, as structural segregation does not necessarily imply enzymatic independence between the two domes.

Consider two representative chains, A and F of the complex belonging to two distinct domes. CYS2699 in Mtb FAS-I (CYS1305 in yeast) is a conserved catalytic residue in KS, while the residue SER1787 (SER180 in yeast) in ACP is the site where the PPT arm binds. [9, 27] Figure 1b shows ACP_A_ stalled at KS_A_ of the same chain in our KS’ model. The PPT arm is pointing far from KS_A_, in a direction that is away from its catalytic residue. In order to validate our modeled conformation of the PPT arm, we measured the distance between ACP_A_ SER1787 C_*α*_ and KS_A_ CYS2699 C_*α*_, which represents the distance that the PPT arm has to extend to at the KS domain during fatty acid synthesis. This distance is measured as 34.32 Å in our model. In experimentally reported SC FAS-I structures, the same distance is measured to be 35.2 Å in PDB ID 6TA1, and 35.67 Å in PDB ID 8PS1. On the other hand, the length of the PPT arm is known to be in the range of 18-21 Å.[12, 26, 37] Thus, with ACP_A_ at KS_A_, its catalytic residue CYS2699 is out of reach of the PPT arm.

Remarkably, upon analyzing multiple chains in the modeled complex, we found that the PPT arm of ACP_A_ is pointing towards CYS2699 of KS_F_ (Figure 1c). The modeled PPT arm conformation nearly perfectly fits into the catalytic pocket of KS_F_. A similar structural observation has been made in Ref. [16] for *T. lanuginosus* FAS-I, where catalytic clefts of KS are found to point towards opposite domes. In fact, careful analysis of the Mtb FAS-I structure with missing ACP reported in Ref. [9] (PDB ID 6GJC), which our model is based on, reveals that KS of each dome is indeed oriented in such a way that its catalytic pocket is facing the opposite dome. Our modeled PPT arm conformation agrees with this “inverted” orientation of KS. The distance between ACP_A_ SER1787 C_*α*_ and KS_F_ CYS2699 C_*α*_ is measured to be 19.18 Å in our model. The same distance is 19.06 Å in SC FAS-I from PDB ID 6TA1. [20] The PPT arm of each chain is within a comfortable distance to extend to the catalytic residue of the KS in the opposite dome through its catalytic pocket. Thus, despite the structural segregation of ACP domains into upper and lower domes, this cross-dome catalytic coupling suggests that the two domes cannot function independently, but instead operate in a functionally dependent manner.

### 3.2 Stability of the complex

The overall stability of the simulated complex was assessed using the root mean square deviation (RMSD) and root mean square fluctuations (RMSF) of each simulation trajectory. Each trajectory was aligned to the initial frame of the production run before calculating these quantities. Thick lines in Figure 2a show the RMSD of C_*α*_ of protein minus the residues corresponding to the mobile PL, ACP and CL, averaged over all six chains as a function of time for all replicas. For all simulated replicas, the RMSD stabilizes below 5 Å in 300 ns. The RMSF of C_*α*_ of the entire protein, averaged over all six chains and all replicas is shown in Figure 2b. ACP and the linker regions exhibit relatively higher fluctuations, which can be attributed to the inherently mobile nature of ACP and the intrinsically disordered character of the linkers. We further examined the structural stability of ACP by calculating the RMSD of its eight helices. Thin lines in Figure 2a show the six-chain-average RMSD as a function of time for all replicas. The analysis indicates that all eight helices of ACP remain stable throughout the simulation in all trajectories, with the RMSD remaining below 3 Å.

**Figure 2.**
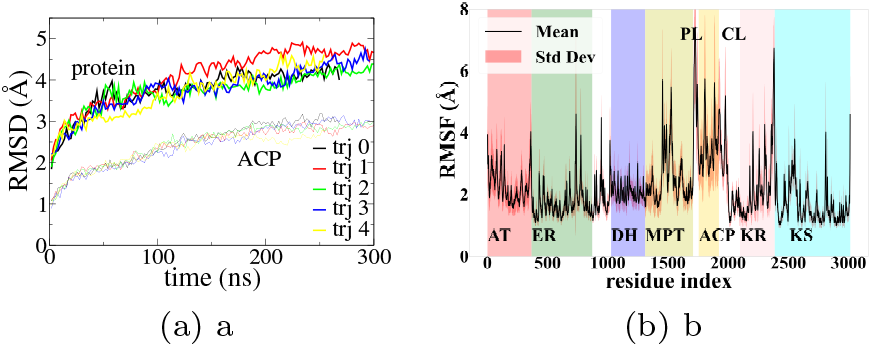
Stability of the Mtb FAS-I complex. (a) Six-chain-average RMSD of C_*α*_ atoms as a function of time in all trajectories. Thick lines shown the RMSD of the protein minus the mobile PL, ACP and CL. Thin lines show the RMSD of the eight helices of ACP. (b) RMSF of all C_*α*_ atoms of the protein averaged over six chains and over all trajectories. Mean RMSF ± standard deviation of each domain is also included.

Stability of ACP at KS was monitored using distance between center-of-mass (COM) of ACP and KS domains in the simulation trajectories. ACP consists of 8 helices, of which 4 helices form the catalytic core that carries out catalysis, and the remaining four form the structural core. The COM distance between ACP’s catalytic core and KS of the same chain in one replica is shown in Figure 3a, while the COM distance between ACP’s structural core and KS of the same chain is shown in Figure 3b. The same two distances with KS of the opposite dome are shown in Figures 3c and 3d, respectively. Lack of any systematic increase in these distances suggests a stable conformation with ACP persistently interacting with the two KS domains throughout the simulation. The catalytic core of ACP maintains a relatively small distance from KS of the opposite dome, which remains stable throughout the simulation trajectory. The distance of either core of ACP from KS of the same chain fluctuates to a greater extent compared to their relatively stable distance from KS of the opposite dome. The same behavior is observed in all other simulation trajectories (data not shown). The nearly constant distances observed in our simulation trajectories suggest a strong interaction between the catalytic core of ACP and the KS of the opposite dome in addition to a relatively weak but stable interaction between both cores of ACP and KS of the same chain.

**Figure 3.**
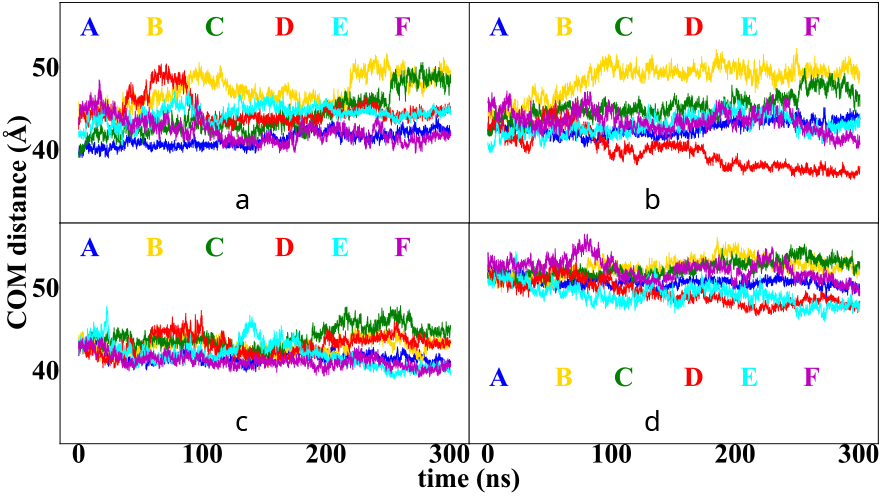
Stability of the conformation with ACP at KS in each chain of replica 1. Time-dependence of the distance between centers-of-mass of (a) the catalytic core of ACP and KS of the same chain, (b) the structural core of ACP and KS of the same chain, (c) the catalytic core of ACP and the cross-dome KS, (d) the structural core of ACP and the cross-dome KS. The alphabets A to F represents the six chains of Mtb FAS-I.

Although catalytic core of ACP has been observed to directly interact with surface residues of other FAS-I domains in fungal species, the structural core of ACP has not been reported to strongly interact with these domains. [40] We have earlier reported surface residues of the catalytic core of ACP that strongly interact with DH residues in Mtb FAS-I, while the structural core of ACP has limited interactions with DH. [24] In contrast, in the conformation with ACP at KS studied here, we observe that the catalytic and structural cores of ACP remain anchored at KS, which contributes to additional structural stabilization of ACP at KS.

### 3.3 Residue-level interactions

As suggested in the preceeding section, ACP seems to persistently interact with surfaces of two KS domains in Mtb FAS-I, one from the same dome and the other from the opposite dome. The structural arrangement observed in our model, where two KS domains provide an extended interaction surface for ACP, is similar to that seen in fungal FAS-I. [20]

The similarity score obtained between SC KS (6TA1, resid 1050-1706) and Mtb KS (6GJC, resid 2501-3091) domain using ClustalW 2.1 Multiple Sequence Alignment is about 29.80%, suggesting a strong homology between KS domains of FAS-I complexes in the two species. The SC KS residues reported to form interaction with ACP are as follows. On the spoke forming parts of KS: residues GLN1123, GLU1124, ILE1126, GLU1128, PHE1134 and PHE1135 (corresponding Mtb KS residues: VAL2518, SER2519, PHE2521, GLU2523, VAL2529 and VAL2530). Near the entrance region of the catalytic pocket of KS: residues GLU1139, ASP1203, GLN1207, ASP1267, ASN1272, ASP1273, MET1346, ARG1416, SER1417, PRO1419 and ALA1420 (corresponding Mtb KS residues: GLU2534, ASP2598, SER2602, GLY2661, ASN2666, ASP2667, MET2740, THR2810, SER2811, PRO2813 and ALA2814). [26] Of these 17 residues, 8 residues, *viz*. GLU1128, ASP1203, ASN1272, ASP1273, MET1346, SER1417, PRO1419 and ALA1420 in SC have corresponding conserved residues GLU2523, ASP2598, ASN2666, ASP2667, MET2740, SER2811, PRO2813 and ALA2814 in Mtb. Further, 5 of the non-conserved residues GLU1124, ILE1126, PHE1134, GLN1207, and ASP1267 in SC correspond to similar residues SER2519, PHE2521, VAL2529, SER2602, and GLY2661 in Mtb. The remaining residues are neither conserved nor similar between SC and Mtb.

In ACP, 16 residues, *viz*. LYS158, LYS161, LYS179, SER180, THR181, GLU185, SER227, ARG230, ARG231, SER234, GLY239, GLY240, THR242, ILE243, THR244 and ARG276 have been reported to interact with KS in SC. [26] Corresponding residues in Mtb ACP are SER1765, MET1768, ALA1786, SER1787, SER1788, GLN1792, ALA1838, ASP1841, GLN1842, THR1845, SER1850, GLY1851, ARG1853, PRO1854, GLY1855 and ALA1901, which includes the two conserved residues SER1787 and GLY1851. Of the remaining 14 non-conserved SC residues, LYS158, THR181, GLU185, SER227, SER230, ARG231, SER234 and GLY239 correspond to similar residues SER1765, SER1788, GLN1792, ALA1838, ASP1841, GLN1842, THR1845 and SER1850 in Mtb. The remaining residues are neither conserved nor similar between SC and Mtb.

Figure 4a shows the percentage frames in which each of the KS residues listed in the preceding paragraph, based on homology with SC, are within a cutoff distance of 4.5 Å from any ACP residue in all chains and in all simulation trajectories. The distance cutoff was selected based on the definition of native contacts used by Best *et al*. (2013) for heavy atoms of the residues. [4] Data for KS residues from the same chain as ACP are shown in red color, while the data for residues from the cross-dome KS are shown in blue color. Residues that maintain interaction for more than 50% of the simulation trajectories (arbitrary cutoff, shown as bright bars) are identified as those having persistent interactions in the following discussion. Nearly half of the 17 KS residues known to be interacting with ACP in SC FAS-I are also seen to persistently interact in our Mtb FAS-I simulation.

**Figure 4.**
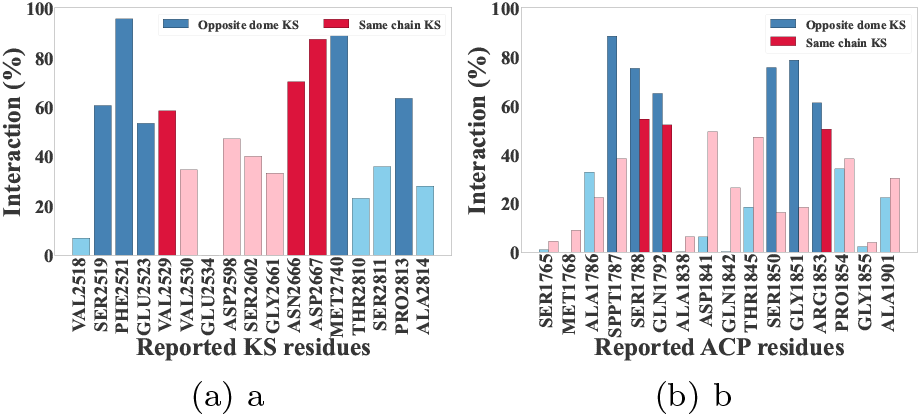
Interaction of the Mtb (a) KS residues of the same chain (red) and opposite dome (blue) with ACP and (b) ACP residues with same chain KS (red) and opposite dome KS (blue) within 4.5 Å across six chains of five trajectories corresponding to the reported for SC KS. the latter provides an additional site for electrostatic interaction with ARG1789.

Similar data for ACP residues listed in the preceding paragraph is shown in Figure 4b. Red color shows data for ACP residues that are within the cutoff distance of KS residues of the same chain, while blue color shows data for ACP residues that are within the cutoff distance of cross-dome KS residues. Nearly half of the ACP residues known to be interacting with KS in SC are also seen to persistently interact in our Mtb FAS-I simulation.

Our analysis also revealed novel residues in Mtb FAS-I that maintain persistent interaction in all chains and all simulation trajectories. These residues have not been reported to interact in SC. [26] Figure 5a shows 11 additional residues belonging to either KS of the same chain (red color) or to the cross-dome KS (blue color) that persistently interact with ACP residues of a given chain. Figure 5b shows 7 additional ACP residues that maintain a persistent interaction with same chain (red color) and cross-dome (blue color) KS.

**Figure 5.**
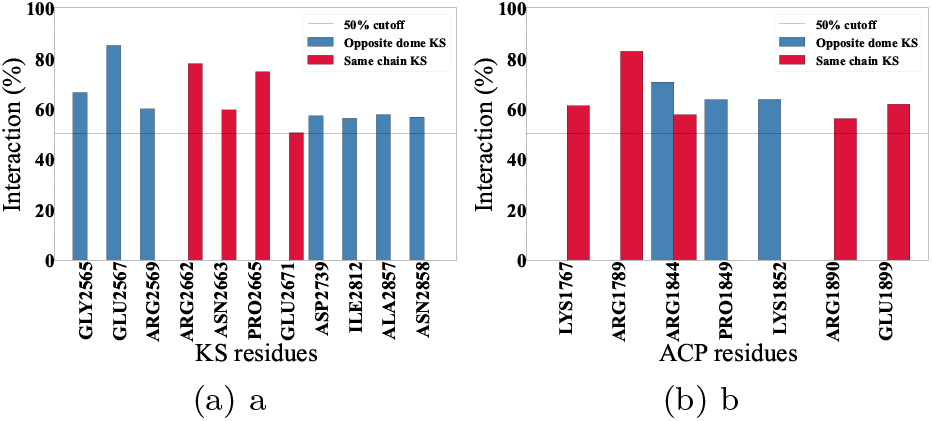
Additional (a) KS residues of same chain (red) and opposite dome (blue) interacting with ACP, (b) ACP residues interacting with the same chain KS (red) and the opposite dome KS (blue) within 4.5 Å and with more than 50% interaction across six chains of five trajectories. Dotted line represents 50% interaction.

Figure 6 shows a heat map of pairs of residues, one from ACP and the other from KS, that are involved in maintaining contacts, as seen in Figures 4 and 5. Color indicates the percentage of simulation frames where a given pair of residues is within the cutoff distance. Conserved residues are marked with an asterisk.

**Figure 6.**
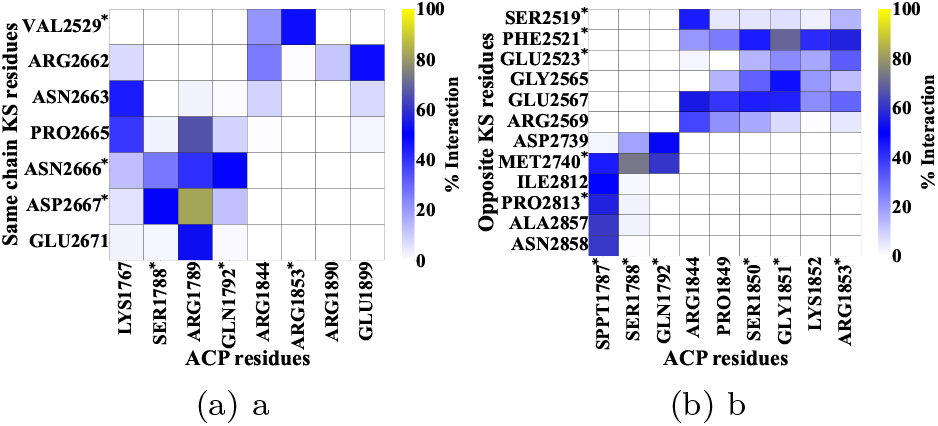
Persistent contacts between ACP and KS residues. Percentage of simulation frames in which is 3.37 Å, which is likely to be within the required distance for the proposed reaction mechanism. [13] ACP residues are within interaction distance from *d*_1_ is below 5 Å, 7.5 Å and 10 Å in 2.9%, 6.6% and residues of (a) KS of same chain and (b) cross-dome KS.

The residue pair ARG1789-ASP2667 is observed to have the most persistent contact in all simulation trajectories, staying within the cutoff distance for 82% of simulation frames. This residue-pair belongs to the same chain, and provides a strong attractive electrostatic interaction between the catalytic core of ACP and KS. ARG1789 also maintains persistent contact with KS residues PRO2665 and GLU2671. Both of these KS residues are in vicinity of ASP2667, and

Among the cross-dome residues, we observe a persistent contact between the residue-pair SER1788-MET2740, which stays within the cutoff distance for 73% of simulation frames. The neighboring SER1787 also shows persistent contact with cross-dome KS residues ILE2812, PRO2813, ALA2857 and ASN2858. Similarly, GLN1892 maintains persistent contacts with ASP2739 and MET2740. However, given the proximity of SER1787 and SER1788 to ARG1789 and the large common surface shared by the two KS domains, these persistent contact may be a result of the strong ARG1989-ASP2667 interaction.

Notably, GLY1851 from the structural core of ACP also maintains a persistent contact with PHE2521 of the cross-dome KS (69.5% frames), while the nearby charged residues ARG1844 and ARG1853 maintain a persistent contact with cross-dome GLU2567 and PHE2521, respectively. Of these, the residue pair ARG1844-GLU2567 (54.9%) provides electrostatic interactions between the structural core of ACP and the cross-dome KS.

These results suggest that ASP2667 is an important binding site for the charged residues in the catalytic core of ACP. Similarly, PHE2521 and GLU2567 are potentially important binding sites for structural core of ACP.

### 3.4 PPT arm dynamics

Conformational dynamics of PPT arms of all six chains in all simulation trajectories was investigated. Of the several conformations observed, we note two peculiar conformations: a fully-extended PPT arm positioned within the catalytic pocket of KS and a “sequestered” PPT arm directed inwards into the ACP catalytic core (Figure 7). Such conformational dynamics of the PPT arm has been reported in bacterial PKSs and fungal FAS-I systems. [19, 35]

**Figure 7.**
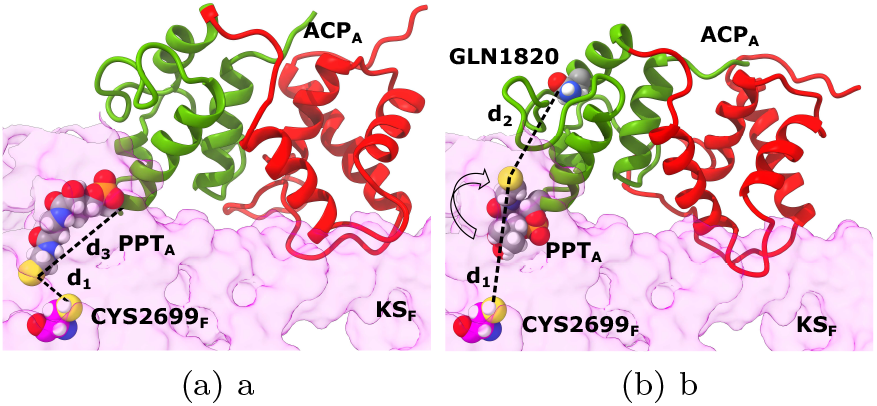
Two distinct conformation of ACP stalled at KS. (a) PPT arm extended conformation buried in KS pocket and (b) PPT arm sequestered conformation directed in ACP pocket.

The fully extended conformation of the PPT arm has also been modeled in SC FAS-I and hFAS structures. [26, 7] Our simulation trajectories of Mtb FAS-I with ACP at KS consist of several frames that correspond to this fully extended conformation of the PPT arm. We characterize this conformation by monitoring the distance *d*_1_ between the S atom of SER1787 and the S atom of CYS2699 in all our simulation trajectories. CYS2699 is a catalytic residue in KS, which is conserved across different species. [23] The minimum value of distance *d*_1_ sampled in our simulation 14.3 % frames, respectively. Thus, the fully extended conformation of the PPT arm reaching into the catalytic pocket of KS is sampled a significant number of times in our simulation with ACP stalled at KS. We also report a strong correlation between the conformations of the PPT arm and the volume of the KS pocket, as seen in Figure 8a.

**Figure 8.**
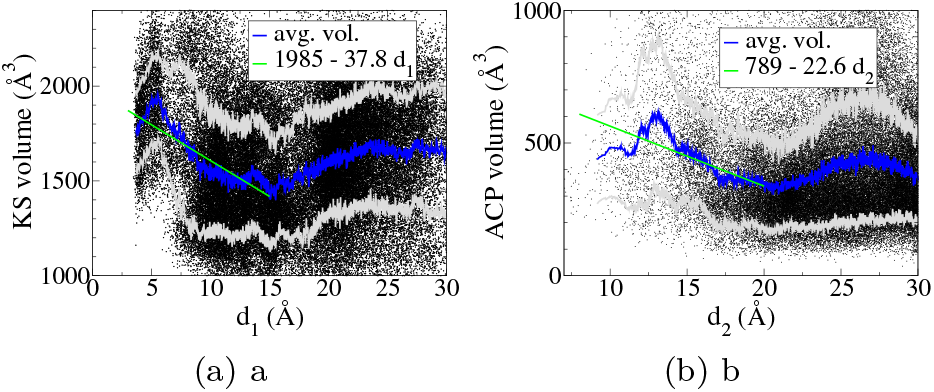
Volume of (a) the KS pocket versus distance *d*_3_ and (b) the sequestering region in ACP versus distance *d*_4_. Lower values of *d*_3_ and *d*_4_ correspond to fully extended and sequestered conformations, respectively. Raw data corresponding to each simulation frame is shown as a black dot. The 200-point running average volume (blue) ± standard deviation (gray) is seen to decrease with increasing distance over short distances. The green lines show a linear fit to the raw data for *d*_3_ *<* 15 Å and *d*_4_ *<* 20 Å, respectively.

Another peculiar conformation is a sequestered PPT arm, which facilitates burial of the growing substrate into ACP. It is speculated that due to physical trapping of ACP in the closed structure of FAS-I, there is no need to sequester the growing substrate in type I systems, unlike in type II systems. [34] Sequestration of the substrate has not been reported in ACP of rat FAS-I. [36] In contrast, it has been observed in NMR spectroscopy experiments of SC FAS-I ACP. [26, 35] In these structures, the substrate is reported to be sequestered in the hydrophobic region between the *α*−3 and *α*−4 helices of ACP, although the sequestered conformation is limited to about 10-25% of the total conformations observed. [35] We characterize this conformation using the distance *d*_2_ between the S atom of SER1787 and the C−*α* atom of a distant residue GLN1820 in the *α* − 4 helix of ACP. Even in the absence of the acyl chain tethered to the S atom of SER1787 in our simulation, we observe a minimum value of *d*_2_ = 6.15 Å. We also sample *d*_2_ *<* 15 Åin ~ 2.7% of the frames, *d*_2_ *<* 18 in ~ 7.9% of the frames and *d*_2_ *<* 20 in ~ 14.0% of the frames. We find a significant correlation between the conformations of the PPT arm and the volume of this region in ACP. The average volume of this region over all simulation trajectories increases with a decrease in the distance *d*_2_, as seen in Figure 8b. These distances throughout the trajectory is shown with respect to time for all six chains in Fig 9 a-c.

**Figure 9.**
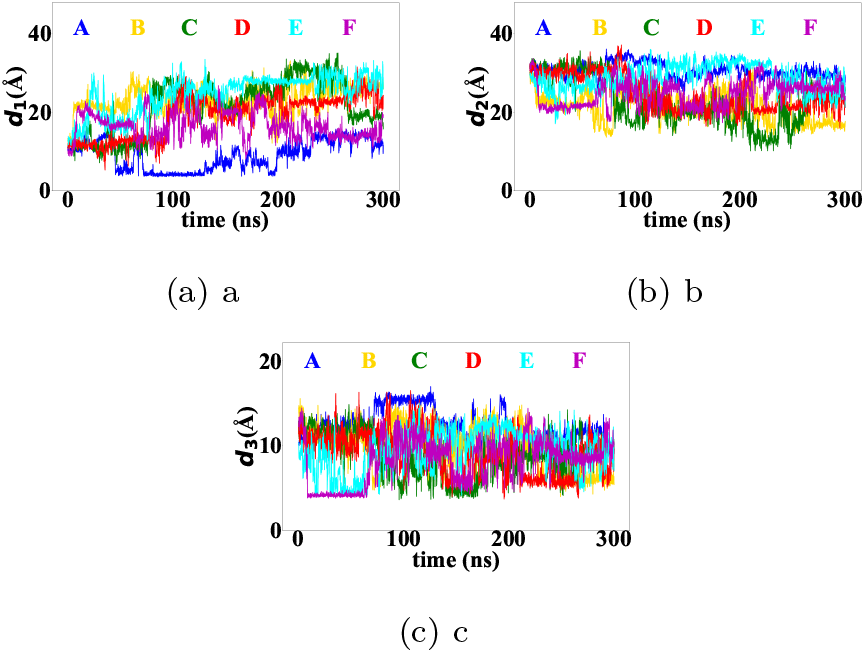
(a) Distance of tip of PPT arm (S of SER1787) and catalytic residue (S of CYS2699) in replica 1. (b) Distance of tip of the PPT arm (S of SER1787) and C*α* GLN1820 on *α* − 4 of catalytic core. (c) End-to-end distance of tip of PPT arm (S of SER1787) and recognition helix (CA of SER1787) in replica 1

### 3.5 Gating mechanism in Mtb FAS-I KS domain

In *de novo* synthesis of fatty acids and polyketides, a phenylalanine residue has been proposed as gate-keeper residue that opens and closes the access to the catalytic site CYS2699 in KS. [31, 30] PHE2958 is indeed observed at the entrance of the cross-dome KS pocket in which our PPT arm conformation is modeled. Its orientation is dynamic in all simulation trajectories. Figure 10a shows a sample simulation snapshot in which this residue is oriented nearly orthogonal to the KS pocket, restricting access to the catalytic residue and resembling a “closed” gate. Figures 10b and 10c show the residue oriented parallel to the pocket, resembling an open gate. The angle *θ* between the C_*α*_ atom of LEU2956, C_*β*_ atom of PHE2958 and C_*z*_ of PHE2958 is used to characterize orientation of this residue. The former two atoms are approximately parallel to the local orientation of the KS catalytic pocket. A histogram of *θ* in all chains and in all simulation trajectories is shown in Figure 11. The histogram clearly indicates the predominance of two “states” that are characterized by the bimodal distribution of *θ*. Sample simulation snapshots corresponding to the two peaks are shown in the inset of Figure 11. No significant correlation is observed between *θ* and the distance *d*_1_, perhaps due to the fact that our simulation did not include an acyl chain. An investigation on the role of the length of acyl chain in the KS step of fatty acid synthesis will be carried out in the future.

**Figure 10.**
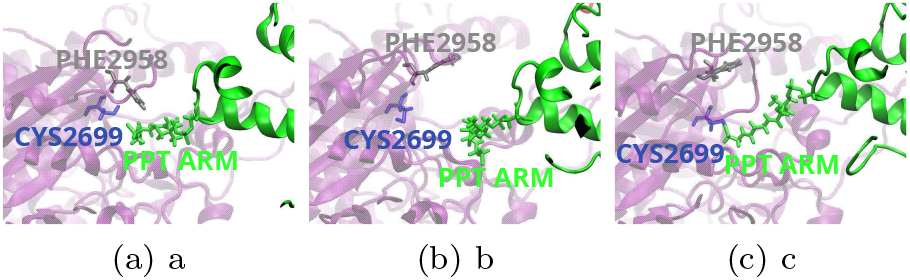
Snapshots showing various conformations of the gating residue PHE2958

**Figure 11.**
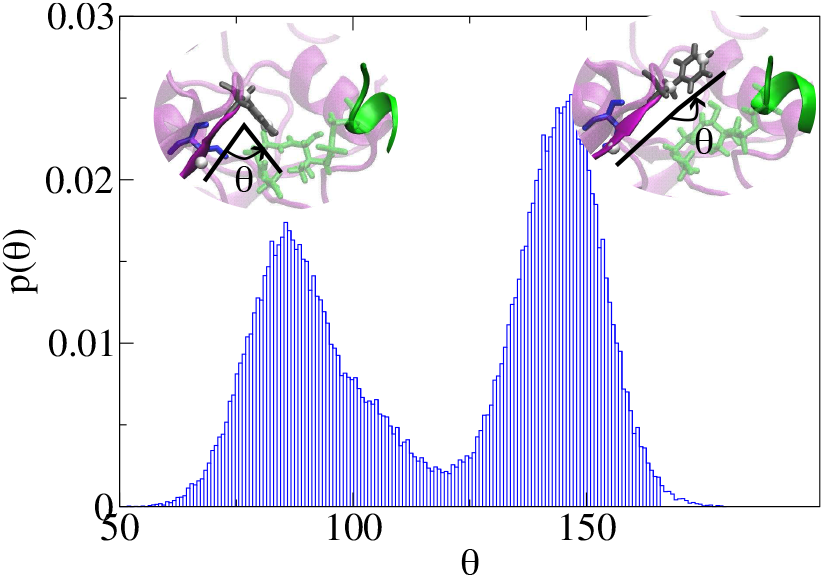
Histogram of the gating residue PHE2958 orientation angle *θ*.

### 3.6 ACP accessibility to condensation enzymes within the dome

It has previously been reported that the yeast ACP can access all catalytic sites depending on the length of the linkers, despite the fact that the ACP and linker structures at the catalytic sites were not resolved [16]. ACP_*A*_ can access KS_*A*_ while interacting with the adjacent MPT_*C*_, which we designated as the KS’ conformation. In contrast, the alternative pathway mediates condensation at a different KS domain (KS_B_) while retaining interaction with the same chain MPT as ACP. We refer to this arrangement as the KS” conformation. Another scenario is when the ACP can access the farthest KS domain (KS_C_), which we designate as KS”‘ conformation. These conformations are shown in Figure 12. In the KS’ conformation, the linker adopts an extended conformation which enables its interactions with neighboring ACPs. In the KS” conformation, the linker assumes a more compact, folded configuration, and does not exhibit any interactions with neighboring ACPs. In contrast, the KS”‘ conformation is characterized by a crisscross-like arrangement of linkers, indicative of increased spatial overlap with potential steric interference.

**Figure 12.**
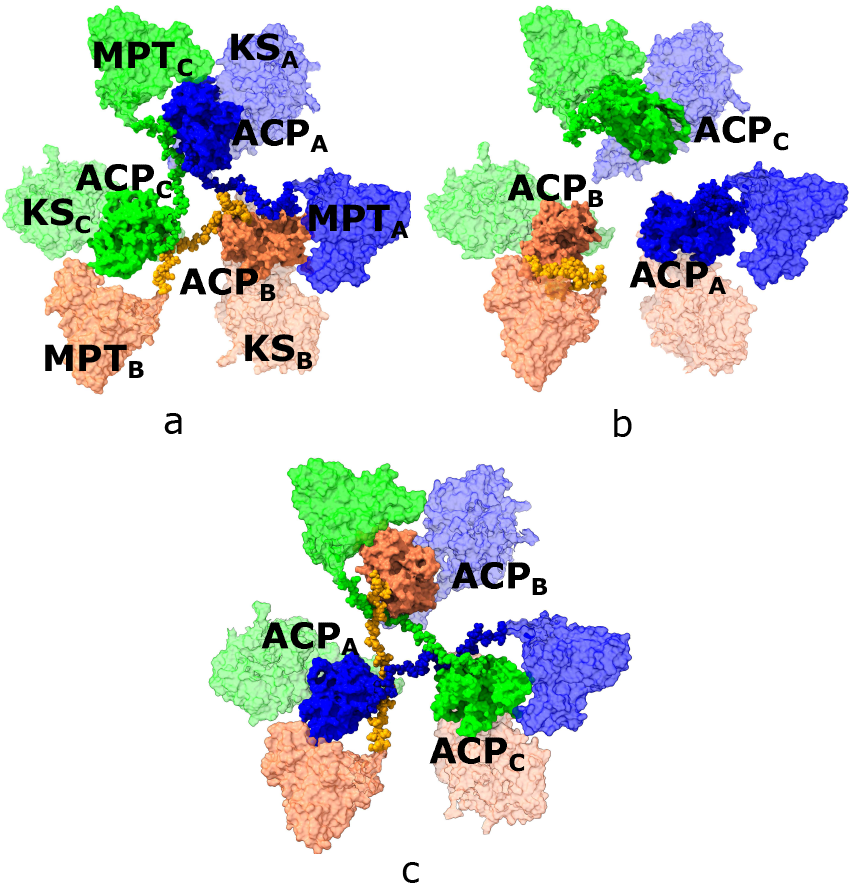
Accessibility of ACP to (a) KS’, (b) KS” and (c) KS”‘ conformations within a dome and conformation of PL.

There is no experimental evidence in support of either of these conformations, due to the difficulty in resolving the structure of the highly flexible linkers, PL and CL. The possibility of these three conformations raises a key mechanistic question: if ACP can access all three conformations, which pathway does it preferentially adopt to mediate fatty acid elongation? To address this, we measure the contour length of the PL linker, measured between its N-terminal residue (1710) and C-terminal residue (1750) in PyMOL. When ACP is in the KS’ conformation, the PL contour length is 77 Å; in the KS” conformation, it is 40.3 Å; and in the KS”‘ conformation, it extends to 132.5 Å. At first glance, these measurements suggest that the KS” conformation may represent the shortest linker-constrained pathway relative to KS’. However, this apparent geometric advantage must be evaluated in the context of ACP’s orientation and the overall shuttling trajectory that involves all other domains. Nonetheless, the KS”‘ conformation appears structurally unfavorable, as it displays the largest PL contour distance, along with a propensity for linker entanglement, as seen from Figure 12. Our analysis suggests that the ACP shuttling mechanism likely follows a pathway in the KS’ conformation, while the alternative pathway involving KS” is also likely. However, a detailed investigation of these conformations is necesssary.

## 4 Conclusion

Our structural and computational analysis aimed to elucidate the evolutionary significance of the hexameric assembly of Mtb FAS-I. Mechanistic insight emerged from the condensation step, highlighting its central role in the ACP shuttling pathway and in coordinating catalysis across the complex. Our results demonstrate that the Mtb FAS-I complex utilizes catalytic domains from both the upper and lower domes during condensation at the KS domain, indicating that the two domes are functionally interdependent during the condensation and elongation steps. This inter-dome coupling has not been explicitly reported previously. Fungal FAS-I, which share structural homology with Mtb FAS-I, is generally presumed to operate through independent domes. To investigate these mechanisms, we modeled previously unresolved structural regions and validated the ACP structure against the recently resolved experimental structure. To our knowledge, this study presents the complete fully functional active Mtb FAS-I model incorporating cofactors and a post-translationally modified ACP with the PPT arm. With ACP positioned at the KS domain, we have characterized ACP interactions with KS domain of both domes. These interactions can guide the mutagenesis study to design inhibitors for fatty acid synthesis. Another crucial detail is ACP’s PPT arm that adopts two distinct conformations within the KS catalytic pocket and ACP sequestering pocket. We found a correlation between these conformations and pocket volumes. The pocket volume increases when the PPT arm is deeply positioned within the pockets. We have also investigated the accessibility of ACP within the reaction chamber. Our analysis suggests that KS’ and KS” are the favourable conformations, while the KS”‘ conformation has a propensity for PL entanglement, provided all ACPs are at the KS domain. A further simulation study is required to support the comment and is backed by the ACP shuttling pathway based on the recently obtained high-resolution structure.

