## Supplemental Information for "Modeling Type-I Fatty Acid Synthase with Acyl Carrier Protein at Ketoacyl Synthase"

May 15, 2026

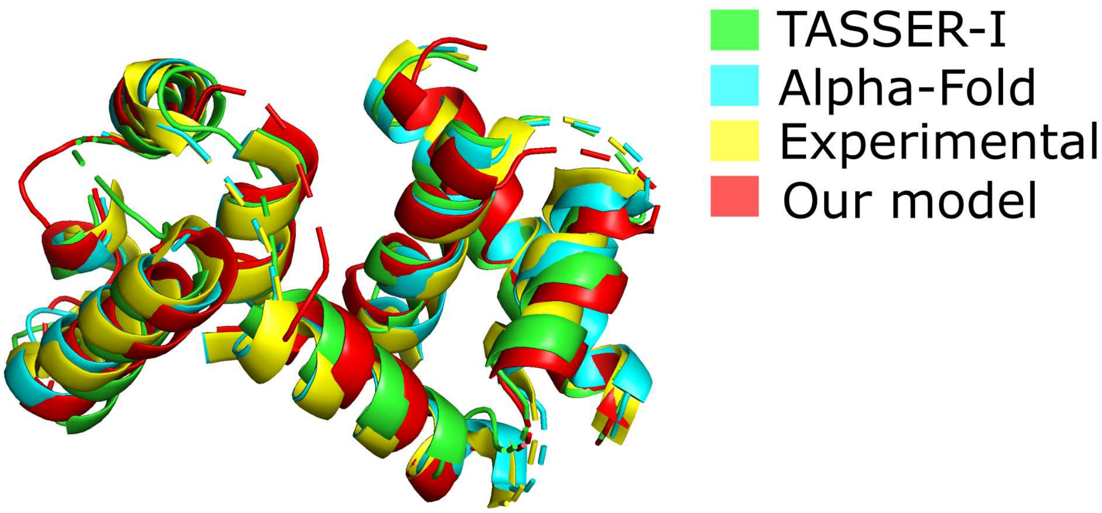


Figure SI-1: Comparison of modelled ACP structure with TASSER, AlphaFold, experimental and our previously reported structure.
